# Effects of Mango–Maize and Mango–Cassava Agroforestry Systems on Arbuscular Mycorrhizal Fungi Communities and Soil Properties in Southern Ethiopia

**DOI:** 10.64898/2026.04.16.718657

**Authors:** Aynalem Gochera Sade, Yonas Ugo Utaile, Bart Muys, Arne Devriese, Olivier Honnay, Margaux Boeraeve

## Abstract

Subsistence agriculture in sub-Saharan Africa faces persistent productivity challenges due to low soil fertility, limited inputs, and increasing climate variability. Agroforestry can offer a sustainable strategy for smallholder systems by enhancing soil quality and the presence of arbuscular mycorrhizal fungi (AMF) in crop roots. Using a canopy-based radial sampling design, we assessed the influence of Mangifera indica (mango) trees on soil properties and AMF communities in maize and cassava in southern Ethiopia. Illumina MiSeq sequencing identified 908 AMF operational taxonomic units (OTUs) from 7 families, dominated by Glomeraceae. While soil properties, including pH, total nitrogen (TN), organic carbon, and potassium, were significantly affected by the distance from mango trunks, this was not the case for AMF community composition and AMF richness and diversity. Host identity, rather than distance from the mango trees, was the primary driver of AMF community composition, with distinct and host-specific assemblages in mango, maize, and cassava roots. Soil nutrients influenced AMF diversity differently across hosts. In maize–mango systems, TN positively affected observed richness (S_obs_) and Shannon diversity (N1), whereas Olsen P negatively affected N1 and Simpson diversity (N2). In cassava–mango systems, TN enhanced S_obs_, and Olsen P positively influenced expected richness (S_exp_). Overall, these findings demonstrate a decoupling between mango-induced soil fertility enhancement and crop AMF community composition and diversity, rather emphasizing the roles of host type and soil nutrients in structuring AMF communities. Without demonstrating direct benefits, we at least show that mango can be effectively integrated into smallholder maize and cassava production without compromising the AMF communities, while enhancing key soil fertility indicators. Maintaining adequate nitrogen levels while avoiding excessive phosphorus inputs may help sustain stable AMF communities in agroforestry systems.

## INTRODUCTION

Subsistence-based agriculture in sub-Saharan Africa (SSA) continues to face persistently low productivity, largely due to limited access to essential agricultural inputs and the growing impacts of climate change (Giller 2020; Shimeles et al. 2018; Watts et al. 2024). As a result, current food production levels remain inadequate to meet the demands of the rapidly growing population (Onyeaka et al. 2024). In addition, unsustainable farming practices have accelerated land degradation, leading to declining soil fertility and reduced agricultural output, thereby intensifying food insecurity and increasing vulnerability to climate-related shocks (Fagbemi et al. 2023; Hossain et al. 2020; Saleem et al. 2024). Addressing these challenges requires sustainable intensification of climate-resilient crop production on existing farmland, alongside efforts to diversify rural livelihoods (Amede et al. 2023; Nguyen et al. 2023; Shilomboleni et al. 2024; Shin et al. 2024).

Agroforestry, which involves the intentional integration of trees with crops and/or livestock in agricultural landscapes, is a promising approach to address the many sustainability challenges faced by smallholder agriculture in SSA (Amadu et al. 2020; Mbow et al. 2014; Sithole & Olorunfemi 2024). Agroforestry diversifies farm products by providing products such as nuts and fruits, contributing to more varied and nutritious diets and increasing resilience to food shortages (García-López et al. 2024; Kuyah et al. 2020). Additionally, it can buffer local microclimates, thereby enhancing the climate resilience of agricultural production systems (Fashing et al. 2022; Ndoli et al. 2017; Zignol et al. 2023). Finally, it helps to combat land degradation by enhancing soil fertility through the addition of organic matter and improving water availability (Kuyah et al. 2019; Marques et al. 2022; Octavia et al. 2023; Ngaba et al. 2024;). For example, observational studies from Ethiopia suggest that agroforestry systems are often associated with improved soil fertility indicators. Parkland agroforestry systems have been reported to exhibit higher soil nitrogen, phosphorus, and organic carbon contents compared to adjacent monoculture fields (Ljalem et al. 2024), while homegarden agroforestry systems have been associated with substantially greater soil organic carbon and phosphorus levels (Wolle et al. 2021). Similar patterns have been reported in other regions of sub-Saharan Africa, where agroforestry practices are frequently linked to enhanced soil nutrient status (Kuyah et al. 2019).

A relatively underexplored mechanism by which agroforestry can enhance soil fertility is by increasing the abundance and diversity of arbuscular mycorrhizal fungi (AMF) in crop roots, either through AMF transfer from tree roots to crop roots, or by improving soil conditions that favor AMF. Belonging to the phylum *Glomeromycota*, AMF are found in association with nearly all crop species and they can play a significant role in enhancing crop yields (Begum et al. 2019; Gebremeskel et al. 2024; Wu et al. 2022). They enhance host plant uptake of nutrients, particularly phosphorus and water (Birhane et al. 2023; Tang et al. 2022), and contribute to improved soil structure (Fall et al. 2022). Although AMF are generally not highly host-specific, certain crop–AMF taxa combinations have demonstrated greater efficiency than others, at least under greenhouse conditions (Van Geel et al. 2016). Overall, AMF benefits can be especially pronounced in low-productivity and low-input agricultural systems, where they may significantly boost productivity and resilience (Rillig et al. 2019; Rog et al. 2025).

Trees in agroforestry systems may support AMF in crops by maintaining active AMF populations that can enhance nutrient exchange and root colonization, thereby improve soil fertility and promoting sustainable agriculture. For example, crop root colonization assessments have shown increased AMF colonization in maize near trees in smallholder farming systems in Malawi, effectively promoting AMF-mediated nitrogen uptake by the crop (Dierks et al. 2022).The effect, however, seems to be strongly dependent on the specific agroforestry tree species (Dierks et al. 2021). A similar AMF morphology-based study in southern Ethiopia found greater AMF colonization and spore density in tree-based systems than in monocrop lands (Masebo et al. 2023). Furthermore, DNA sequencing-based evidence has indicated that tree presence can alter AMF community composition, promote AMF diversity, and network complexity in wheat and coffee roots, possibly contributing to higher yields in agroforestry systems (Broeckhoven et al. 2025; Qiao et al. 2025). Nevertheless, how crop AMF diversity depends on specific tree vs. crop species combinations in low input smallholder agroforestry systems remain underexplored, highlighting the need for more targeted, species-specific studies.

In Ethiopia, maize (*Zea mays*) remains a cornerstone of food security (Abate et al. 2015), while cassava (*Manihot esculenta*) is emerging as a versatile dietary staple, traditionally consumed boiled and increasingly integrated into *injera,* a fermented flatbread that serves as the country’s primary staple food, through flour blending with teff (Bogale et al., 2022; Tafesse et al., 2021). Despite their significance, the productivity of both crops is constrained by declining soil fertility and moisture stress (Assefa et al., 2020). Although fertilizer applications can enhance crop yields, their use among smallholder farmers is often limited by access and affordability, while concerns over the environmental impacts of synthetic fertilizers underscore the need for sustainable nutrient management strategies (Markos et al. 2023; Sinta & Dansa 2023; Tafesse et al. 2021).

In this study, we explored how single standing *M. indica* (mango) trees affect soil properties and AMF richness, diversity and community composition in maize and cassava roots within low-input parkland agroforestry systems of the Southern Ethiopian Rift Valley. Using Illumina high-throughput sequencing on 183 maize, 180 cassava and 29 mango root samples, we profiled AMF community composition and diversity associated with these two vital crops along a distance gradient from single standing mango trees. Specifically, the objectives of this study were (i) to assess how distance from mango trees influences selected soil properties, (ii) to evaluate the effects of mango trees on AMF community composition and richness and diversity in maize and cassava roots, (iii) to compare AMF community composition and diversity among maize, cassava, and mango roots and (iv) to determine how selected soil properties influence AMF taxon richness and diversity in maize and cassava grown within mango-based agroforestry systems.

## MATERIALS AND METHODS

### Study site and sampling methods

The study area is in Ethiopia’s southern rift valley lowlands on the western shores of Lake Abaya, which is situated 15 km north of the town of Arba Minch (Fig.1). The area has a semi-humid to semi-arid subtropical climate with 800–1200 mm of annual rainfall, with heavy rains falling from March to May and brief rains falling from September through November. The average annual lowest temperature is 17°C and the highest is 30°C (Tomass et al. 2021). Farmers grow maize as a staple crop during the short-wet cropping season, which extends between August to December (Alemu, 2016; Gebre et al., 2020) whereas March to May is the optimal planting period for cassava (Bogale et al., 2022). The soil types of the catchment in which our study sites located are mainly vertisols, luvisols, leptosols, and cambisols with parent material of alluvial fans and lacustrine deposits (Endale et al., 2023).

**Figure 1.**
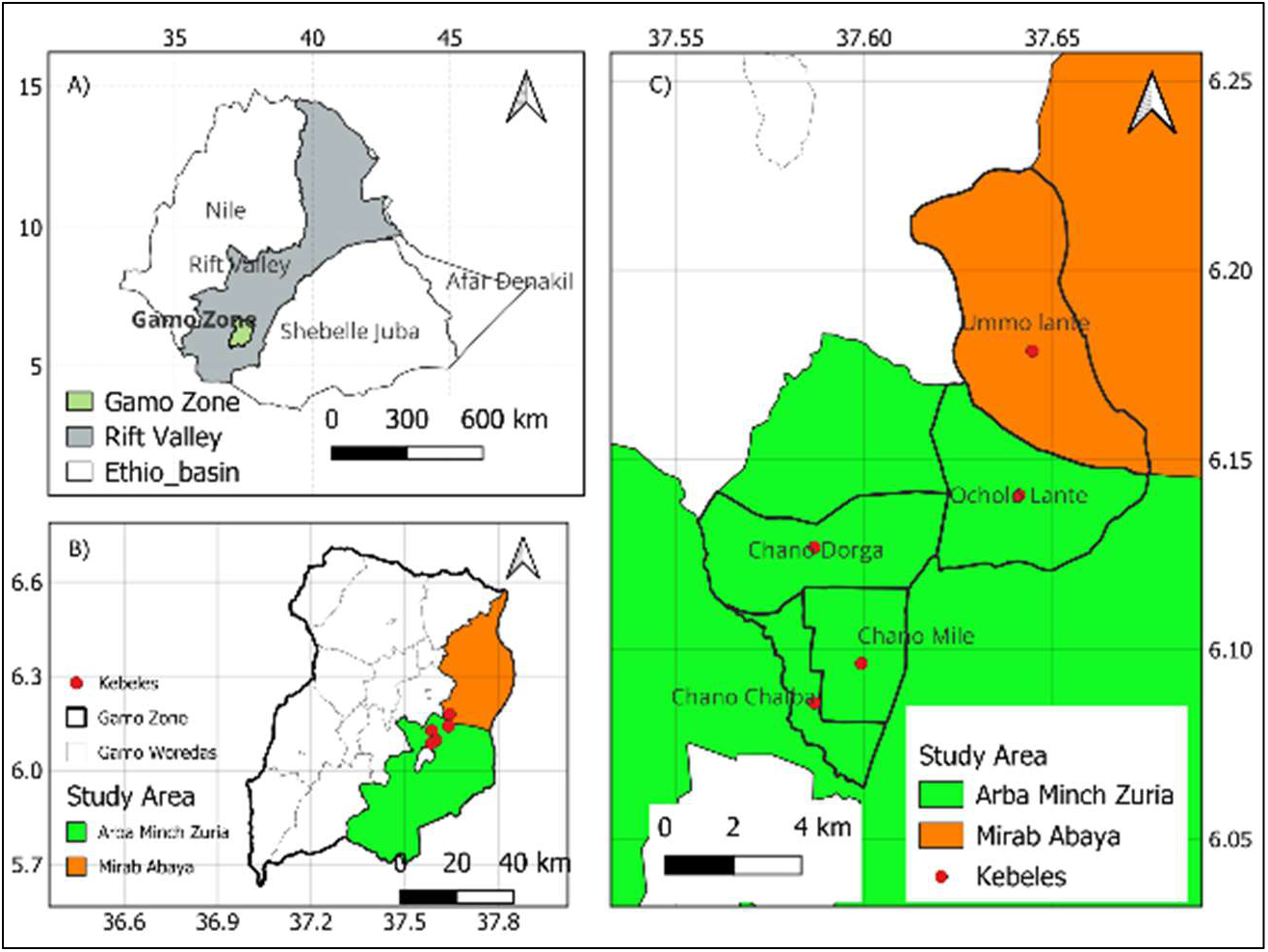
Study area in Ethiopia’s southern rift valley lowlands on the western shores of Lake Abaya.

Fifteen single-standing mature mango trees were selected from farmers’ fields growing cassava, and another 15 from fields growing maize. To minimize external influences, selected trees had to be located at least 40 meters away from each other and from other trees or perennial crops such as banana plants, fences, or dense vegetation (Supplementary Fig. 1).

Soil sampling was conducted extending outward from the trunk at four distances: one-third of the canopy radius, two-thirds of the radius, and four-thirds of the radius (Fig. 2). Additional control samples were taken at the distance of three times the canopy radius, representing open field conditions. At each distance, three soil samples were taken randomly at different positions around the tree to a depth of 0–20 cm using an auger. A total of 360 soil samples were collected and transported to the laboratory for analysis of selected physicochemical properties.

**Figure 2.**
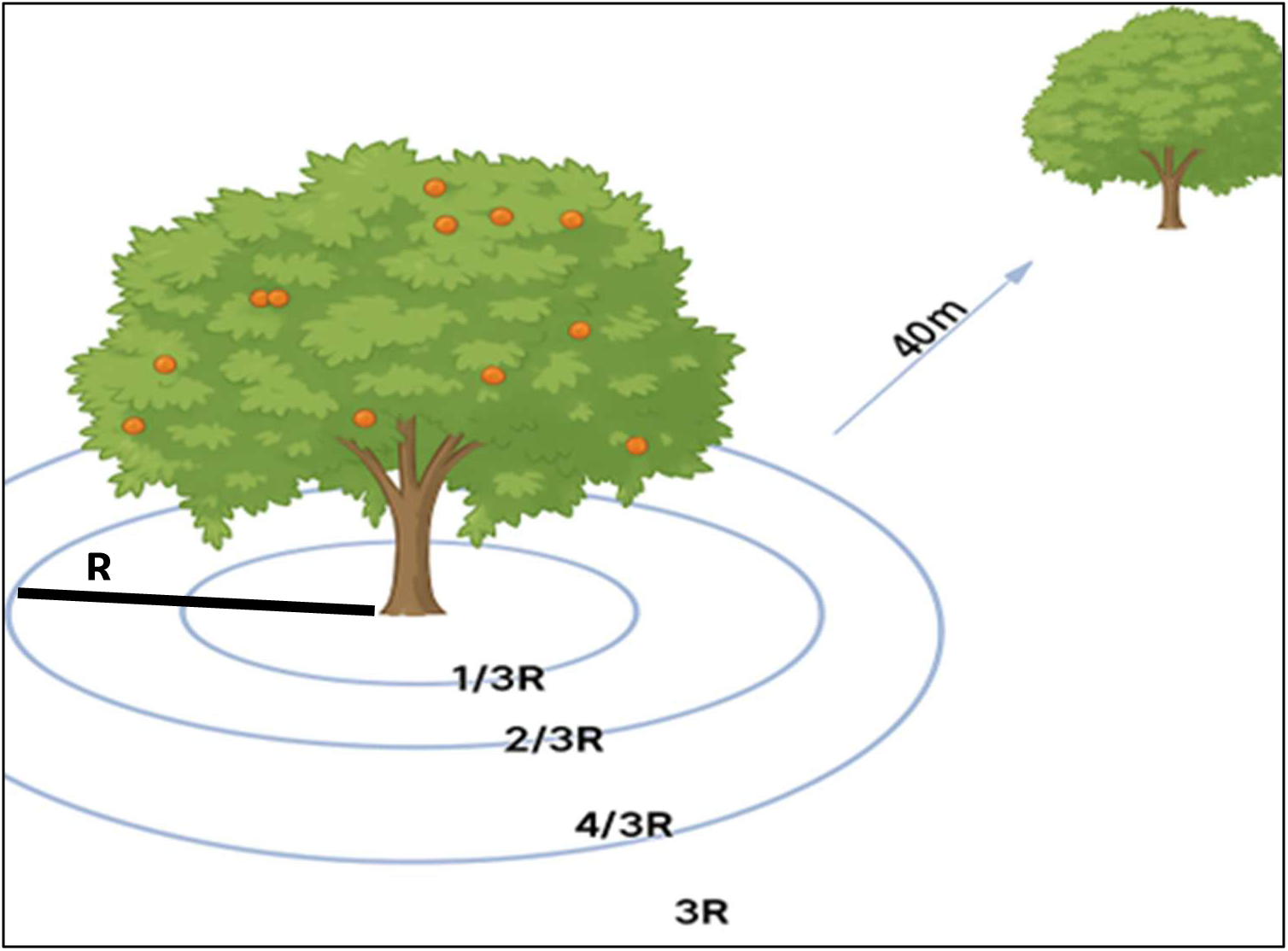
Schematic representation of the soil and root sampling design around mango tree canopies. Samples were collected at distances of one-third (^1^/3R), two-thirds (^2^/3R), and four-thirds (^4^/3R) of the canopy radius (R). A control sample was taken at 3R from the tree base. Selected trees were located at least 40 meters away from other trees.

At each soil sampling distance, one maize or cassava plant was selected, and fine roots were carefully excavated for AMF analysis, resulting in 180 maize and 180 cassava root samples. In addition, fine roots were collected from each mango tree, yielding a total of 30 mango root samples. All root samples were stored in silica gel and transported to the laboratory for DNA extraction.

### Soil Analysis

Soil pH, electrical conductivity (EC), organic carbon (OC), total nitrogen (TN), available phosphorus (Olsen P), and available potassium (K_av_) were determined using standard laboratory procedures from ISRIC (2002) and GLOSOLAN (2019/2021). Soil pH was measured in a 1:2.5 soil-to-water suspension using a digital pH meter (Hanna, Model HI99121, USA), while EC was determined in a 1:5 soil-to-water suspension using an EC meter (Hanna, Model HI99300, USA). Organic carbon content was quantified using the Walkley-Black wet oxidation method. Total nitrogen was assessed via the Kjeldahl digestion method, employing a Kjeldahl Distillation Unit (Model S1, SN-8021268, Behr Labor-Technik, Germany). Available phosphorus was extracted using the Olsen method and determined calorimetrically through the molybdenum blue procedure with an RS-295 spectrophotometer. Available potassium was extracted using ammonium acetate (pH 7.0) and quantified using a flame photometer (Model CL-378).

### PCR amplification and Illumina MiSeq sequencing

Genomic DNA was extracted from the dried roots using the Ultra Clean Plant DNA Isolation Kit (MoBio Laboratories Inc., Solana Beach, CA, USA) following the manufacturer’s instructions. Extracted DNA was diluted with PCR-grade water prior to amplification. The AMV4.5NF/AMDGR primer pair was used for PCR, as it selectively amplifies AMF community sequences targeting the highly variable region of the small subunit (SSU) rRNA gene (Garo et al., 2022; Sato et al., 2005; Van Geel et al., 2014). PCR reactions were performed on a BioRad T100 thermal cycler (Bio-Rad Laboratories, CA, USA) in a 20 µl reaction volume containing 0.15 mM of each dNTP, 0.5 µM of each primer, 1× Titanium Taq PCR buffer, 1 U Titanium Taq DNA polymerase (Clontech Laboratories, Palo Alto, CA, USA), and 1 µl of genomic DNA. The thermal cycling conditions were as follows: initial denaturation at 94 °C for 2 minutes; 35 cycles of 94 °C for 45 seconds, 58 °C for 45 seconds, and 72 °C for 45 seconds; followed by a final extension at 72 °C for 10 minutes. Following PCR, the concentration of the amplified products was measured using a Qubit fluorimeter with the Qubit dsDNA HS assay kit, after purification with Agencourt AMPure XP beads (Beckman Coulter Life Sciences, Indianapolis, IN, USA). DNA fragments of the appropriate size (approximately 350 bp) were excised from the gel and purified using the QIAquick Gel Extraction Kit (QIAGEN). Sequencing was conducted on an Illumina MiSeq platform using the v2 500-cycle reagent kit (Illumina, San Diego, CA, USA) at the Genomics Core Facility (Leuven, Belgium).

### Bioinformatics

The sequence data was processed through the USEARCH sequence analysis tool, following the recommended pipeline (Edgar, 2013). Forward and reverse reads were merged and subsequently oriented using the MaarjAM database (Öpik et al. 2010). Then, reads shorter than 200 bp or with an expected number of errors of more than 0.5 were removed. Sequences were clustered into operational taxonomic units (OTUs), defined at 97% sequence similarity, chimeric sequences were removed, and an OTU table was constructed. Per sample, we filtered out OTUs that were represented by less than 0.01% of the sequences in that sample, to avoid possible erroneous sequences produced during PCR or sequencing (Alberdi et al. 2018). Consensus sequences of the OTUs were BLASTed against the MaarjAM database and unidentified or unsurely identified OTUs (sequence similarity < 95%, query coverage < 90% or e-value ˃ 1e ^−50^) were BLASTed against the NCBI GenBank. Finally, the OTU table was filtered to only contain OTUs identified as belonging to the *Glomeromycota*, which was used in further statistical analyses.

## Statistical Analysis

All statistical analyses were performed using R (version 4.3.2) within the RStudio environment. Prior to analysis, soil variables were examined for skewness and subjected to transformations where necessary to improve normality. Normality was assessed using the Shapiro–Wilk test, and the homogeneity of variance of model residuals was evaluated using Levene’s test and visual inspection of residual plots.

To evaluate the effect of distance from mango trees on individual soil properties, linear mixed-effects models (LMMs) were fitted using the *lmer* function from the *lme4* package (Bates et al., 2015), in combination with the *lmerTest* package (Kuznetsova et al. 2017). LMMs were used to account for the hierarchical structure of the data and the non-independence of observations taken at multiple distances around the same tree. In these models, distance was treated as a fixed effect and tree identity as a random effect. Estimated marginal means were compared to post hoc using the *emmeans* function to identify significant pairwise differences between distances. Principal Component Analysis (PCA) with varimax rotation was employed to explore multivariate relationships among soil variables, while Permutational Multivariate Analysis of Variance (PERMANOVA), implemented via adonis2 in the *vegan* package (Oksanen et al., 2025) was used to test for overall differences in soil characteristics across distances from the trees.

Alpha diversity of arbuscular mycorrhizal fungi (AMF) was assessed using multiple indices, including observed OTU richness (S_obs_), expected richness (S_exp_), Shannon diversity (Hill number N1), and the Simpson diversity (Hill number N2). These metrics were computed using the *estimate_richness* function from the *phyloseq* package (McMurdie & Holmes 2013). Generalized Linear Mixed Models (GLMM) were then used to analyze their response, with tree identity included as a random effect, and both host type (maize, cassava, mango) and distance as fixed effects. Count-based metrics were modeled using a negative binomial distribution via *glmer.nb* from the lme4 package (Bates et al. 2015), while continuous metrics were fitted using the *glmmTMB* function.

AMF community composition was evaluated using Non-Metric Multidimensional Scaling (NMDS) based on Bray–Curtis dissimilarities, implemented with *metaMDS* in the *vegan* package. PERMANOVA was then used to test for significant compositional differences between each crop and mango, as well as across distances from mango trees for both maize and cassava. To identify AMF OTUs specifically associated with each host type, we also performed an Indicator Species Analysis using the *indicspecies* package. The *IndVa*l method (*multipatt* function) was applied to the OTU table using host type as the grouping variable. Statistical significance was evaluated using 999 permutations, and OTUs with p < 0.05 were considered indicator taxa.

To investigate the effect of soil properties on AMF diversity, the diversity indices were analyzed for maize–mango and cassava–mango systems separately. Soil variables were included as predictors in Generalized Linear Models (GLMs) with a log link function. Model selection was performed using the corrected Akaike Information Criterion (AICc) via the *dredge function* from the *MuMIn package* (Bartoń 2023), and the most parsimonious model was used for inference and visualization. Post hoc comparisons of estimated marginal means were conducted using *emmeans*.

## RESULTS

### Effect of *M. indica* trees on soil properties

Soil pH, EC, TN, K_av_ and OC varied significantly with distance from the tree base (Fig. 3). In contrast, Olsen P did not differ significantly across distances (F = 1.37, *p* > 0.05; Supplementary Table 1). Soil pH was significantly higher at a tree base (^1^/3R) compared to all other distances, while pH levels at ^2^/3R, ^4^/3R, and 3R did not differ significantly (Fig. 3). EC showed a declining trend with increasing distance from the tree base (F = 32.7, *p* < 0.001; Supplementary Table 1). EC levels at ^1^/3R and ^2^/3R were significantly higher than those at ^4^/3R and 3R. However, no significant differences were detected between ^1^/3R and ^2^/3R, or between ^4^/3R and 3R (Fig. 3). OC (F = 57.47, *p* < 0.001), K_av_ (F = 117.05, *p* < 0.001) and TN (F = 26.4, *p* < 0.001) all varied significantly with distance from the tree base (Supplementary Table 1). For each of these soil properties, concentrations were highest at ^1^/3R and significantly greater than at all other distances (*p* < 0.001). The ^2^/3R also differed significantly from both ^4^/3R and 3R (*p* < 0.05), whereas no significant differences were observed between ^4^/3R and 3R (Fig. 3).

**Figure 3.**
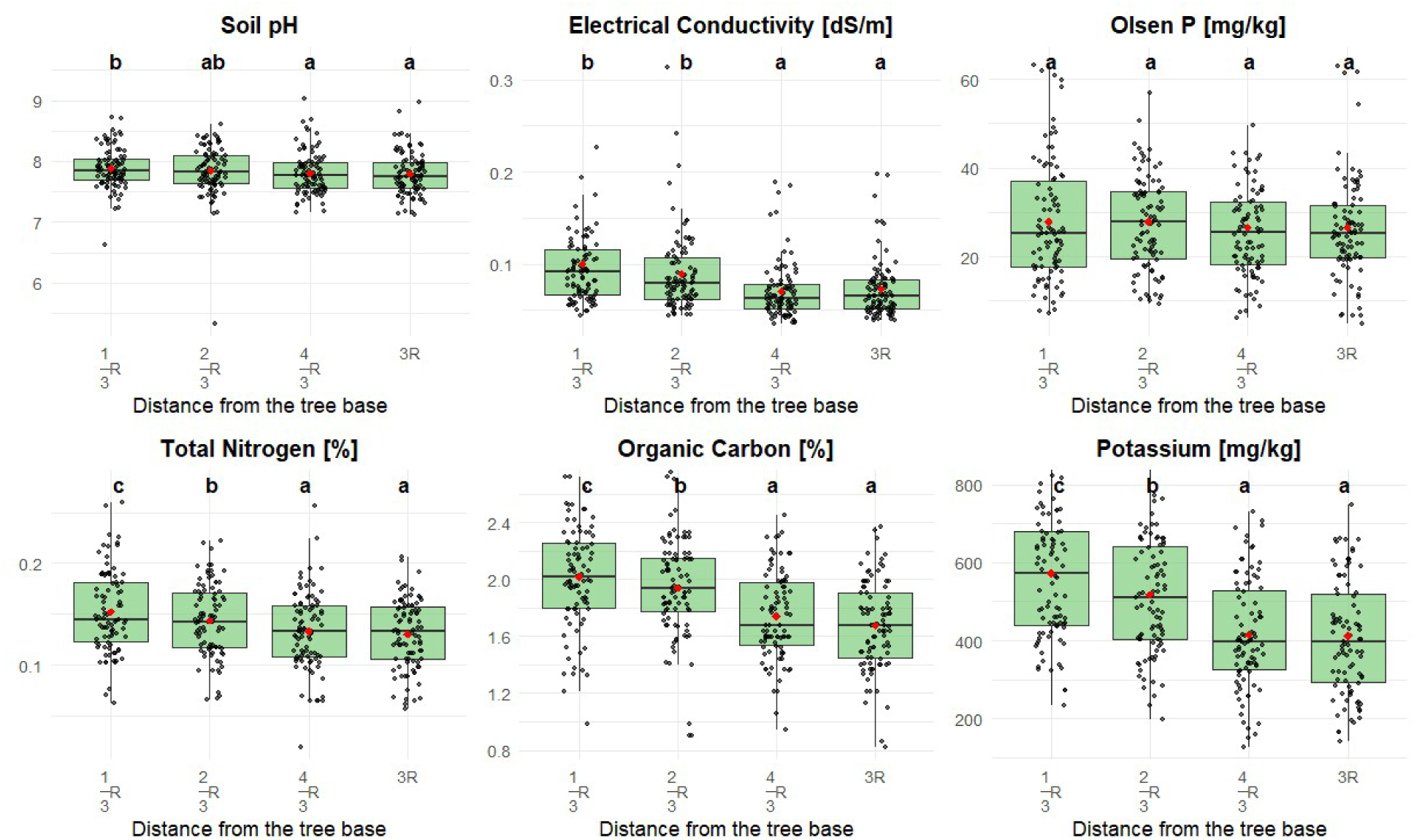
Effects of distance from *mango* trees on selected soil properties in mango-based agroforestry systems. Boxplots display soil properties across four distances relative to the tree base: ^1^/3R, ^2^/3R, ^4^/3R, and 3R, with R the canopy radius. Red dots represent group means; black dots show individual observations. Different lowercase letters indicate statistically significant differences among distances for each soil variable, based on post hoc comparisons.

PCA indicated that variation in selected soil properties across distances from mango tree bases was primarily explained by the first two components (RC1 = 42.1%, RC2 = 26.3%) (Fig. 4). K_av_, OC, TN, and Olsen P showed strong positive loadings on RC1, while electrical conductivity (EC) and pH were more aligned with RC2. A spatial pattern was observed across distances. Samples taken close to the trunk were more concentrated at higher RC1 values, whereas samples taken far from the trunk were located at low RC1 values (Fig 4). PERMANOVA results showed that the multivariate composition of selected soil properties was significantly affected by distance from *M. indica* trees (R² = 0.16, F = 22.05, *p* = 0.001; Supplementary Table 2).

**Figure 4:**
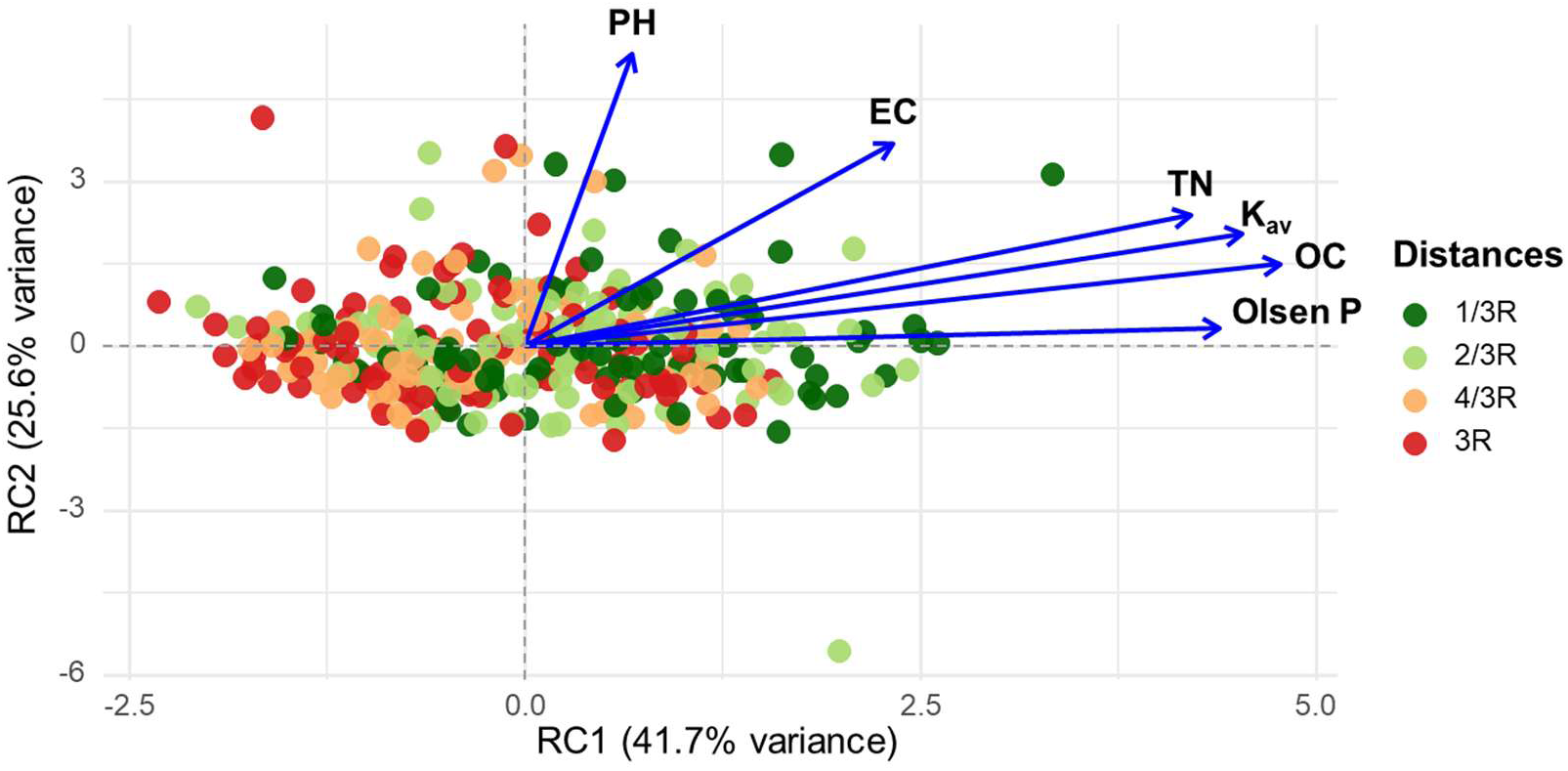
PCA biplot of soil samples collected in mango agroforestry systems at four distances from the mango tree base: 1/3R , 2/3R , 4/3R and 3R , where R is the canopy radius. Arrows indicate the loadings of soil chemical properties. Rotated components RC1 and RC2 explained 42.1% and 26.3% of the total variance, respectively.

### Arbuscular Mycorrhizal Fungi Communities

In total, 170 maize, 157 cassava, and 29 mango root samples were successfully processed and included in the sequencing workflow. Two Illumina MiSeq sequencing runs together generated 10 956 367 sequences belonging to 908 AMF OTUs following quality filtering. Taxonomic classification assigned these OTUs to 9 genera and 7 families. The majority were affiliated with the family *Glomeraceae* (786 OTUs, 94.2% of the sequences), followed by C*laroideoglomeraceae* (3.09%) and *Diversisporaceae* (1.73%), while the remaining families each contributed less than 1% of the total sequences (Fig. 5). At the genus level, Glomus overwhelmingly dominated the community, representing more than 94% of the sequences (Supplementary Fig. 2). Smaller proportions were attributed to *Claroideoglomus* (3.09%), *Diversispora* (1.72%), and *Acaulospora* (0.57%), with all other genera collectively contributing < 0.5% of the total sequences.

**Figure 5:**
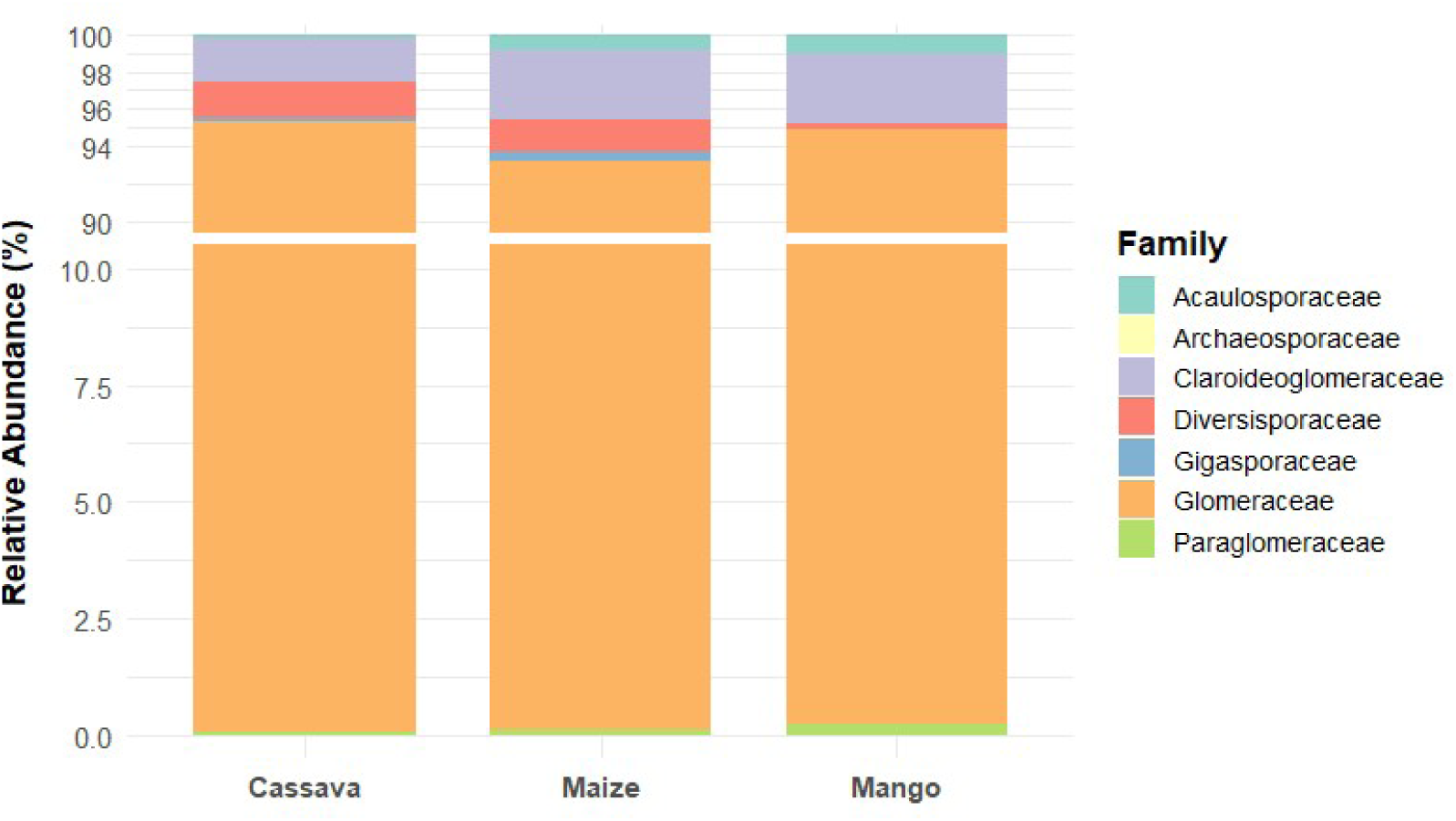
Relative abundance (i.e. proportion of sequences) of AMF families associated with cassava, maize, and mango roots in mango-base agroforestry systems.

Among all identified OTUs, cassava harbored 122 unique OTUs, maize 195, and mango only 8. A substantial number of OTUs (386) were shared between cassava and maize, while far fewer were shared between cassava and mango (2) or between maize and mango (15). Notably, 180 OTUs were common to all three hosts (Supplementary Fig.3). Of the total observed OTUs, 108 (11.9%) were identified as indicator species significantly associated with one or more host types (cassava, maize, or mango). The distribution of AMF indicator families varied among hosts (Supplementary Table 3). *Glomeraceae* was consistently dominant across all hosts, indicating a core AMF group. *Diversisporaceae* was more abundant in cassava and maize than in mango, while *Acaulosporaceae* was more prominent in mango, suggesting host-specific preferences. *Gigasporaceae* was absent from mango, further highlighting compositional differences in AMF communities among the host types.

### Effects of host type and distance on AMF richness and diversity

AMF diversity varied significantly among host types across all diversity indices (Fig. 6; Supplementary Table 4). Observed richness was strongly influenced by host type (S_obs_; χ² = 72.52, p < 0.001), and post-hoc comparisons showed that all three hosts differed significantly from one another. For expected richness (S_exp_; χ² = 36.53, p < 0.001), maize and cassava did not differ significantly, whereas both crops exhibited higher richness than mango. A similar pattern was observed for Shannon diversity (N1; χ² = 24.52, p < 0.001) and Simpson diversity (N2; χ² = 19.52, p < 0.001), where maize and cassava showed comparable diversity levels and both differed significantly from mango. In contrast, no significant differences in AMF diversity were detected across distances from *M. indica* trees for either maize or cassava.

**Figure 6:**
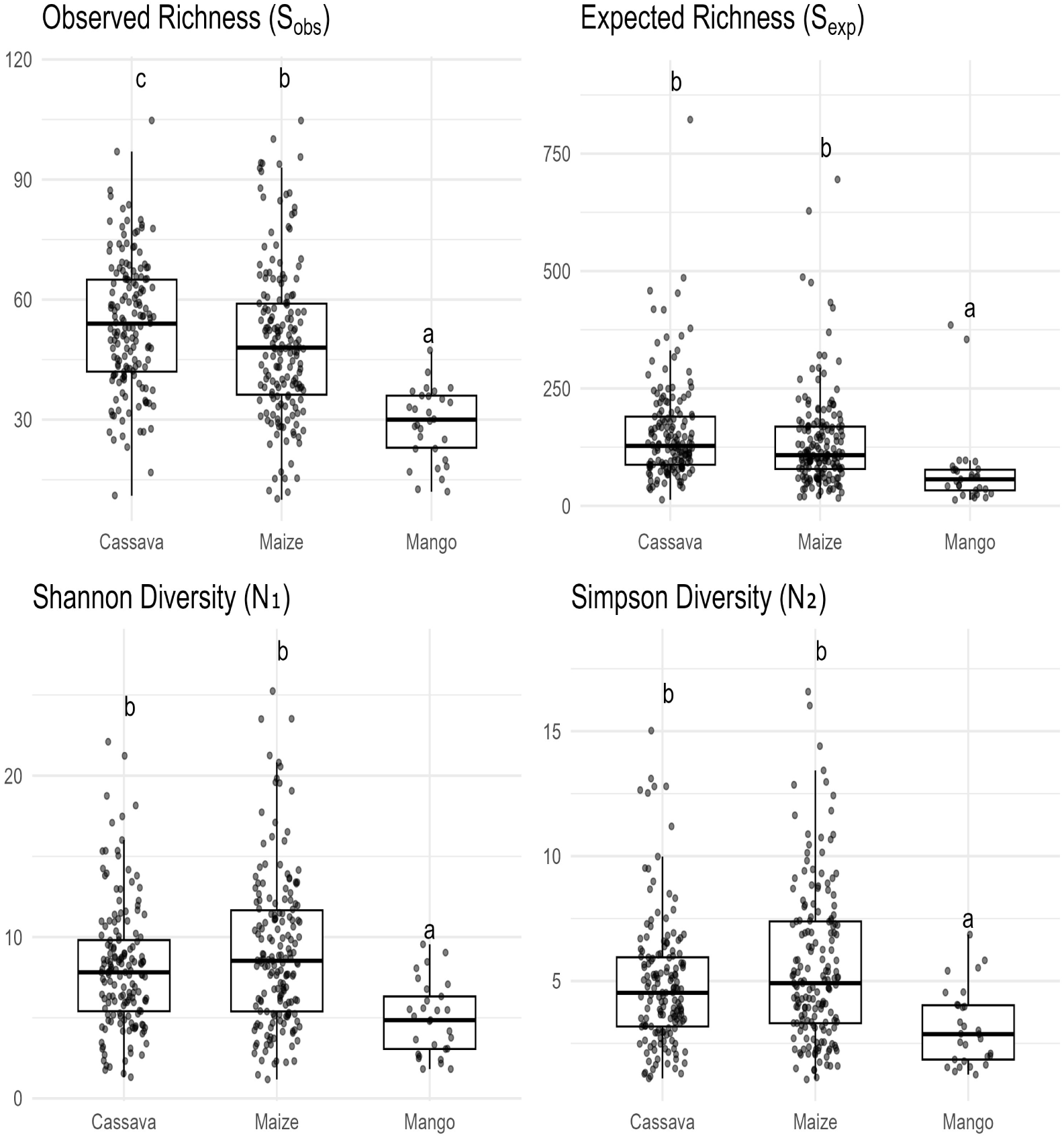
Richness and diversity of AMF across Cassava, Maize and Mango in mango-based agroforestry systems. Different letters above denote statistically significant differences among host types (*p* < 0.05), while the same letter indicates no significant difference based on post-hoc pairwise comparisons.

### Effects of *mango* trees on AMF community composition

PERMANOVA revealed no significant distance-related variation in AMF community composition in either maize (F = 0.893, R² = 0.02, p = 0.67; Supplementary Table *5*) or cassava (F = 0.56, R² = 0.01, p = 0.10; Supplementary Table 6; Supplementary Fig. 4&5). In contrast, host type had a significant effect on AMF community composition (F = 11.91, R² = 0.06, p = 0.0001; Supplementary Table 7). Pairwise comparisons showed the dissimilarity between cassava and mango (R² = 0.11, p < 0.001), followed by maize and mango (R² = 0.09, p < 0.001), with a weak yet significant difference between maize and cassava (R² = 0.006, p = 0.025) (Supplementary Table 8). These patterns are visually supported by the NMDS ordination, which shows clear clustering of AMF communities by host type, particularly separating mango from both cassava and maize (Fig. 7).

**Figure 7:**
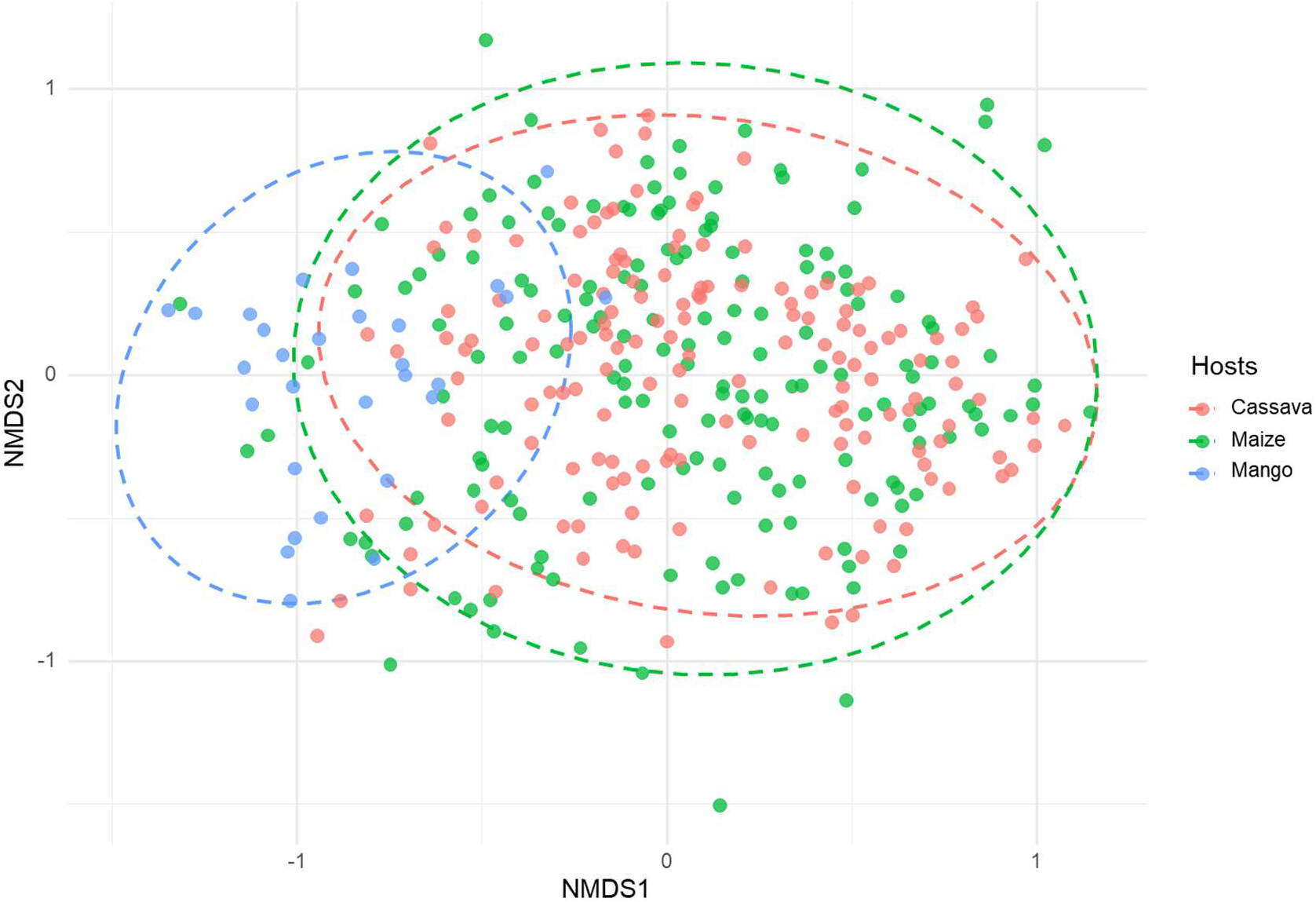
NMDS ordination showing AMF community composition across host types in mango-based agroforestry systems. Points represent individual samples from cassava, maize, and mango roots. Ellipses indicate 95% confidence intervals for each host group, revealing partial separation of AMF communities by host type.

### Effects of soil properties on AMF richness and diversity in maize and cassava roots

Based on AICc model selection, GLMMs identified soil properties that explained variation in AMF diversity metrics for both maize–mango and cassava–mango systems separately. In the maize–mango system, TN had a significantly positive effect on observed richness (S_obs_, p < 0.05) and Shannon diversity (N1, p < 0.05), and a marginally positive effect on Simpson diversity (N2, p < 0.1). In contrast, Olsen P showed a significantly negative effect on Shannon diversity (N1, p < 0.05) and a marginally significant negative effect on Simpson diversity (N2, p < 0.1) (Fig.8, Supplementary Table 9). In the cassava–mango system (Supplementary Fig. 6), TN had a significantly positive effect on observed richness (S_obs_, p < 0.05). Olsen P showed a positive, marginally significant effect on expected richness (S_exp_, p < 0.1) (Fig. 8, Supplementary Table 9).

**Figure 8:**
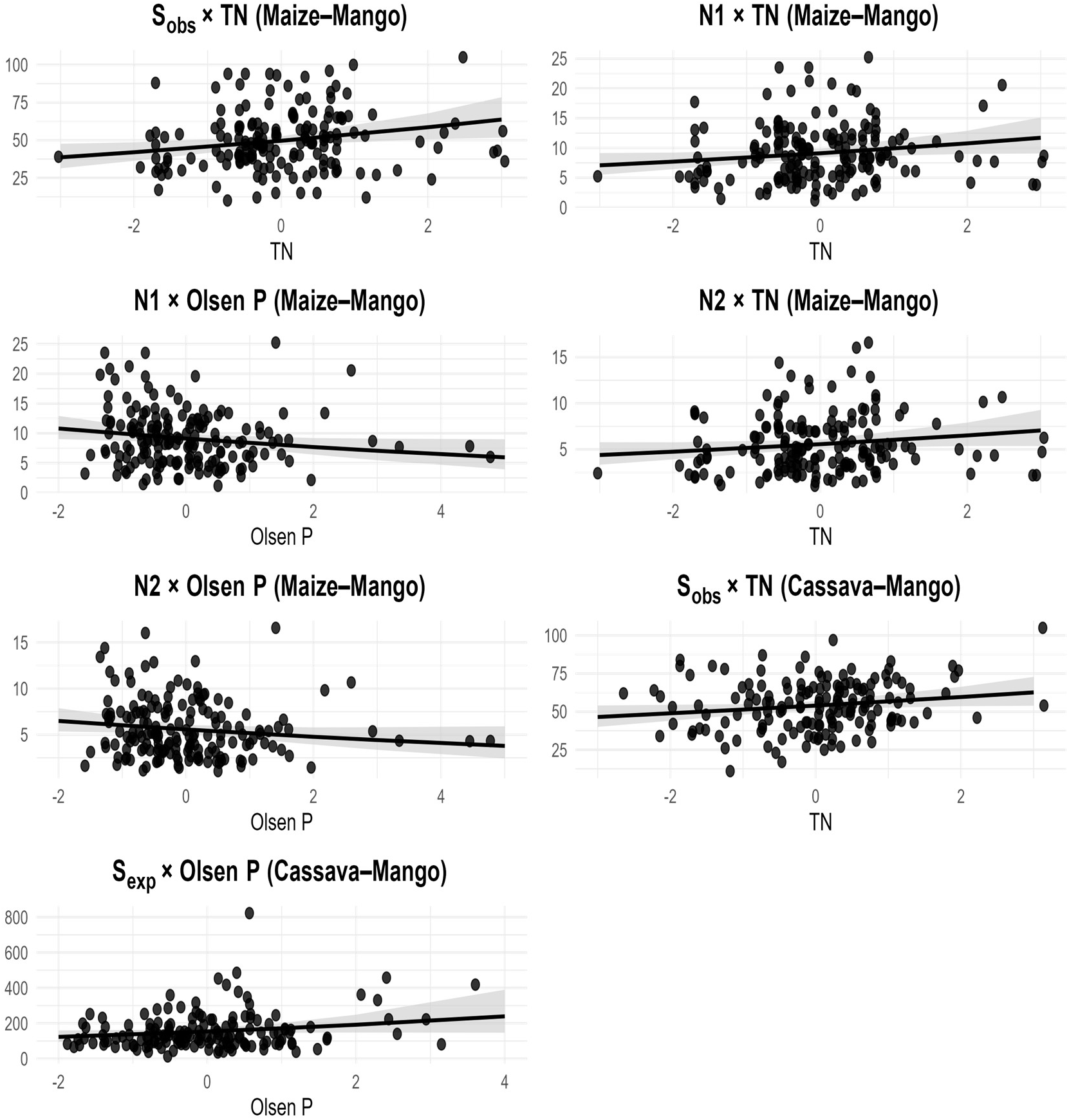
Effects of soil properties on AMF richness and diversity in maize–mango and cassava–mango systems. Lines are fitted GLM predictions (Gamma–log link) with 95% confidence; points represent individual samples. S_obs_: observed richness, S_exp_: expected richness, N1: Shannon diversity, N2: Simpson diversity, TN: total nitrogen.

## DISCUSSION

In this study, we addressed a key knowledge gap regarding the impacts of scattered trees in cassava and maize agroforestry systems in Ethiopia on soil properties, AMF community composition, richness and diversity, as well as the influence of soil properties on AMF richness and diversity. Therefore, we investigated how distances from mango trees affected both soil characteristics and AMF communities. Our main findings are that whereas distance from the tree base significantly influenced selected soil properties, we found no evidence that the presence of mango trees increases AMF diversity or alters AMF community composition in neighboring crops. Contrary to our expectations mango tree roots supported less diverse AMF communities that were distinct from those of maize or cassava roots.

### Effect of Distance from *M. indica* Trees on Soil Properties

The presence of mango trees in agricultural fields positively influences soil quality, particularly in areas close to the tree base. Except for soil pH, which was only higher near the tree trunk, electrical conductivity, potassium, total nitrogen, and organic carbon all exhibited clear decreasing trends with increasing distance from the tree base. These patterns suggest that mango trees enhance soil fertility in comparison to open field conditions or monoculture. Similar spatial patterns have been reported in previous studies investigating scattered trees in parkland agroforestry systems. For instance, Etafa (2022) reported that total nitrogen, available phosphorus, exchangeable potassium, and pH were significantly higher beneath the canopies of different shade trees compared to open areas in the Sayyo District of western Ethiopia. Likewise, Ljalem et al. (2024) reported that both physical and chemical soil properties improved significantly under *Anogeissus leiocarpa* (African birch) and *Stereospermum kunthianum* in the dryland agroforestry parklands of Tigray. Similarly, Gebrewahid et al. (2019) observed that scattered trees such as *Olea europaea* and *Dalbergia melanoxylon* enhanced soil fertility on smallholder farms in the semi-arid regions of Ethiopia.

Despite the well-documented soil-enriching effects of agroforestry tree species, relatively limited research has explicitly examined the role of *M. indica*. Existing studies nonetheless indicate that mango trees improve soil quality across diverse agroecological contexts. Mango-based agroforestry systems have been shown to increase soil organic carbon, organic matter, and nitrogen, reduce bulk density, and enhance soil aggregation and nutrient retention (Nyamangara et al. 2009; Rodrigues et al. 2019; Haider et al. 2024; Dierks et al.202). Improvements in soil pH and macronutrient availability have also been reported in South Asia (Rana 2022). These positive effects are largely attributed to continuous organic matter inputs from mango litter (Pascal & Karim 2024), enhanced nutrient cycling, and microclimatic moderation through canopy shading, which together promote microbial activity, moisture retention, and long-term soil fertility. Participatory assessments further confirm the perceived importance of mango trees in sustaining soil-related ecosystem services in smallholder systems (Gochera et al. 2025).

The improved soil quality observed near mango trees can be driven by interacting mechanisms, including continuous organic matter inputs from leaf litter and fine-root turnover, enhanced microbial activity under shaded and moisture-conserving microclimates, and more efficient nutrient retention through improved soil aggregation (Pascal & Karim 2024). The accumulation of organic matter enhances cation exchange capacity and nutrient buffering, while improved aggregation promotes carbon stabilization and reduces nutrient losses. Together, these processes explain the positive soil responses observed in our study and demonstrate the multifunctional role of *M. indica* in agroecological intensification, sustainable land management, and long-term soil restoration in comparable agroforestry systems.

### AMF Communities in Mango–Maize–Cassava Agroforestry Systems

We found that the AMF communities were dominated by the family *Glomeraceae*, with the genus *Glomus* accounting for over 94% of the OTUs, followed by *Claroideoglomeraceae* (3.09%), represented mainly by *Claroideoglomus*. The dominance of *Glomus* aligns with previous studies reporting its broad ecological distribution and high root colonization efficiency across tropical and subtropical agroecosystems. For example, *Glomus* has been identified as the prevailing genus in mango rhizosphere soils, with relative abundances up to 86% (Yang et al. 2025). Similarly, Mohandas (2012) documented its predominance across several mango rootstocks, attributing this to efficient root colonization and enhanced nutrient uptake and plant vigor. Comparable patterns have also been reported in cassava-based systems in southern Nigeria, where *Glomeraceae* was the most abundant AMF family (Thanni et al. 2022). In coffee agroecosystems, both within and beyond the native range of *Coffea arabica*, *Glomus* species were consistently identified as the most frequently occurring taxa (Bertolini et al. 2018; Broeckhoven et al. 2025). Likewise, Garo et al. (2022) reported a clear dominance of *Glomus* in *Enset* roots in the Southern Ethiopian Rift Valley, followed by members of the *Claroideoglomeraceae* family. Overall, the dominance of *Glomeraceae* likely reflects their broad ecological tolerance, rapid root colonization, and resilience to soil disturbance. These traits enable *Glomus* species to thrive in managed agroecosystems, potentially playing a key role in nutrient cycling and plant–soil interactions.

### Effect of distance and host type on AMF community composition and diversity

Our study showed that the AMF communities associated with cassava, maize, and mango were significantly different, with *Diversisporaceae* contributing most to cassava and maize, and *Acaulosporaceae* contributing most to mango, according to the IndVal analysis. This finding aligns with previous research indicating that host species act as one of the primary ecological filters shaping AMF community structures (Davison et al. 2011; Frew et al. 2025; Öpik et al. 2010; Vázquez-Santos et al. 2025). For example, Geoffroy et al. (2017) found that the composition of AMF colonizing taro (*Colocasia esculenta* L.) roots differed markedly from those associated with *Pterocarpus* species in a swamp forest in Guadeloupe. Similarly, Ilyas et al. (2023) reported that in high-organic-matter soils in Ontario, Canada, crop species was the dominant factor influencing AMF communities colonizing onion and carrot roots. Their findings support the hypothesis that even when grown in the same ecological zone, different host plants can harbor distinct AMF communities.

In our study, mango, a perennial woody crop, supported a more distinct AMF assemblage compared to the annual crops, cassava and maize, which both exhibited greater overlap in community composition. Similar distinctions between perennial and annual hosts have been reported elsewhere (Alguacil et al. 2012; Muleta et al. 2008), and could be due to variation in root architecture, life history traits, and root exudate chemistry. Our results underscore the importance of host plant identity, and especially host plant functional group, in shaping root-associated fungal communities. Importantly, this distinct AMF assemblage associated with mango and the annual crops provide further support of the limited impact the presence of mango has on the AMF communities of those crops in an agroforestry system. The limited colonization of cassava and maize by mango-associated AMF would prevent enrichment or increased AMF richness near the tree trunk, indicating that host specificity may constrain tree-induced enhancement of AMF diversity in mango–maize and mango–cassava agroforestry systems.

The large overlap in AMF community composition between cassava and maize likely reflects their shared functional traits, similar management conditions, and co-cultivation in intercropping systems. In such systems, maize is often interplanted in cassava fields during early growth stages, leading to comparable soil management and fertilization regimes. These similarities may favor the recruitment of generalist AMF taxa, producing more similar communities than those associated with mango.

*The observed AMF richness (S_obs_)* followed the trend cassava > maize > mango, indicating that plant traits, growth habits and management regimes are key factors in structuring AMF communities. Cassava exhibited the highest richness, likely due to its starch-rich, tuberous root system, which offers extensive and nutrient-rich microhabitats favorable to AMF colonization (Thanni et al. 2022, 2024). Maize, an annual cereal crop, showed intermediate richness, which may reflect its moderately fibrous root system and intensive cultivation practices that influence soil microbial dynamics and promote colonization by generalist AMF taxa. In contrast, mango, a perennial woody species, supported the lowest AMF richness. Its long-lived root system may foster more stable but less diverse fungal communities, potentially due to the dominance of a few AMF taxa that competitively exclude others over time. The stable yet less diverse AMF association in mango has also been reported by Yang et al. (2025). Both Shannon (N1) and Simpson (N2) diversity indices were significantly higher in cassava and maize than in mango, indicating greater AMF richness and evenness in the annual crops. Conversely, the lower diversity observed in mango suggests a community structure dominated by fewer, persistent AMF taxa. This pattern may be again attributed to mango’s perennial, woody growth habit and long-lived root system, which tend to support more stable but less dynamic fungal assemblages over time. The reduced AMF diversity in mango may also be linked to allelopathic compounds in its root exudates, which can suppress colonization by fungal taxa, favoring the persistence of more resistant species (Kato-Noguchi & Kurniadie 2020; Aman et al. 2023).

### Effects of Soil Properties on AMF Richness and diversity

We examined how soil properties influence AMF richness and diversity in maize–mango and cassava–mango agroforestry systems. In both the maize-mango and cassava-mango system, AMF diversity was positively correlated with soil N. This aligns with previous findings where moderate nitrogen enrichment enhances AMF colonization (Lenoir et al. 2016; Wang et al. 2024). While nitrogen had a consistently positive effect, the response to phosphorus varied between systems. In the maize–mango system, Olsen P showed a negative association with AMF diversity, reducing Shannon and Simpson diversity. In the cassava-mango system, Olsen P did not have a clear effect on AMF diversity, with a marginally non-significant, positive correlation between S_exp_ and Olsen P and no significant correlations between Olsen P and the other diversity indices. In contrast, in the cassava–mango system, Olsen P was positively associated with S_exp_, indicating a different response pattern. Conversely, the negative relationship between Olsen P and AMF diversity supports previous evidence that high phosphorus availability suppresses mycorrhizal symbiosis, as plants reduce carbon allocation to fungal partners when phosphorus demand is met (Van Geel et al. 2017). However, in the cassava–mango system, the positive effect of Olsen P levels on AMF richness may reflect its role in stimulating fungal activity within the plant–soil interaction zone. This finding suggests the importance of balanced nutrient availability and organic matter in shaping belowground fungal diversity in mango-based agroforestry systems. The positive effect of phosphorus observed in our study contrasts with reports that high soil P suppresses AMF richness (Geel et al. 2017). This discrepancy reflects the context-dependent nature of AMF responses to nutrients. For instance, Broeckhoven et al. (2025) found higher AMF diversity under elevated P in wild forest coffee systems, likely due to host adaptations and site-specific conditions. Our results align with (Qin et al. 2020), who showed that moderate P levels enhance AMF diversity. Collectively, these findings suggest that moderate rather than extreme P availability supports diverse and functionally robust AMF communities in cassava –mango systems. This may relate to cassava’s longer root lifespan and greater mycorrhizal dependency. Additionally, management differences may contribute to maize is often fertilized with di-ammonium phosphate (DAP), reducing mycorrhizal reliance, whereas cassava is typically unfertilized, maintaining nutrient levels favorable for AMF diversity. Overall, these results highlight the interactive role of soil nutrients and management in structuring AMF communities across mango-based agroforestry systems.

## Conclusion

This study demonstrates that the presence of mango trees is associated with improved soil pH, organic carbon, total nitrogen, and potassium, supporting soil fertility maintenance and long-term sustainability in smallholder agroforestry systems, while AMF richness and diversity remain largely unaffected by tree canopy presence. Although we show no beneficial effects, integrating mango trees with maize and cassava at least does not negatively affect arbuscular mycorrhizal fungal communities, indicating compatibility between mango trees and staple food crops. However, AMF community composition differed among host plants, with cassava and maize supporting greater richness and diversity than mango.

Relationships between soil properties and AMF diversity were system-specific. Total nitrogen was consistently associated with increased AMF richness and diversity, whereas phosphorus was negatively associated with diversity in maize–mango systems and showed a marginal positive effect on richness in cassava–mango systems. These patterns demonstrate the importance of balanced nutrient management for sustaining soil microbial communities in agroforestry systems. In mango-based agroforestry systems, maintaining adequate nitrogen levels while avoiding excessive phosphorus inputs may help sustain stable AMF communities and overall agroecosystem functioning.

Overall, *Mangifera indica* can be effectively integrated into smallholder maize and cassava production without compromising AMF communities, while enhancing key soil fertility indicators. Management practices should prioritize adequate nitrogen supply and avoid excessive phosphorus inputs. Future research should explore how mango and other agroforestry species, such as *Kigelia africana* and *Cordia africana*, influence broader ecosystem functions to inform sustainable smallholder agroforestry development in the Ethiopian Rift Valley.

## Acknowledgements

We gratefully acknowledge VLIR-UOS for funding this research. We thank Gerrit Peeters for his outstanding laboratory work, and the late Dr Simon Shibru (Assoc. Prof.) for supervising the field activities. We are also deeply grateful to our field team and to the farmers in Arba Minch Zuria and Mirab Abaya woredas who allowed us to conduct research on their farms.

---

**Figure 6:**
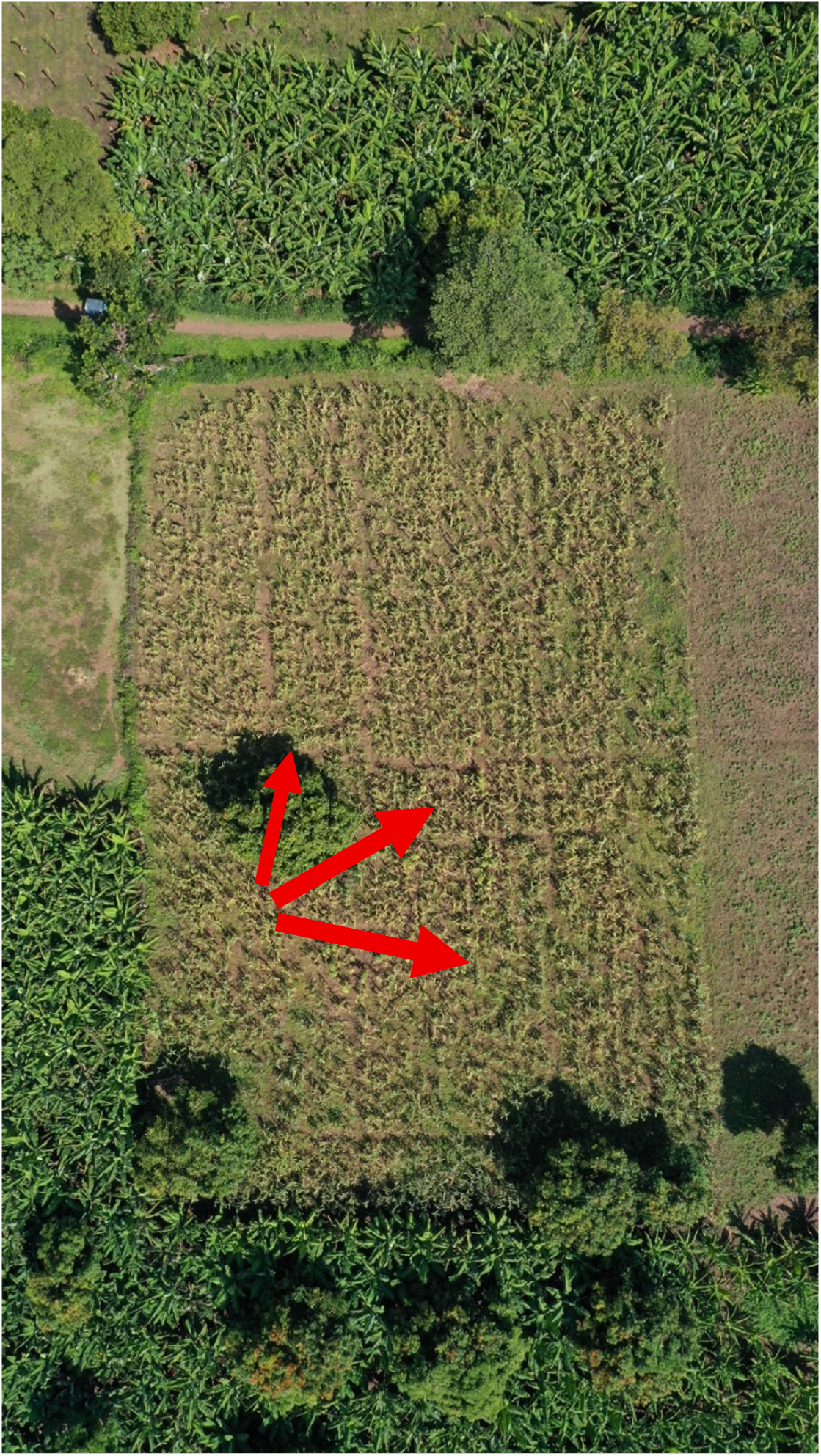
One of the experimental Mangifera indica trees intercropped with maize in Chano Mille.

**Figure 7:**
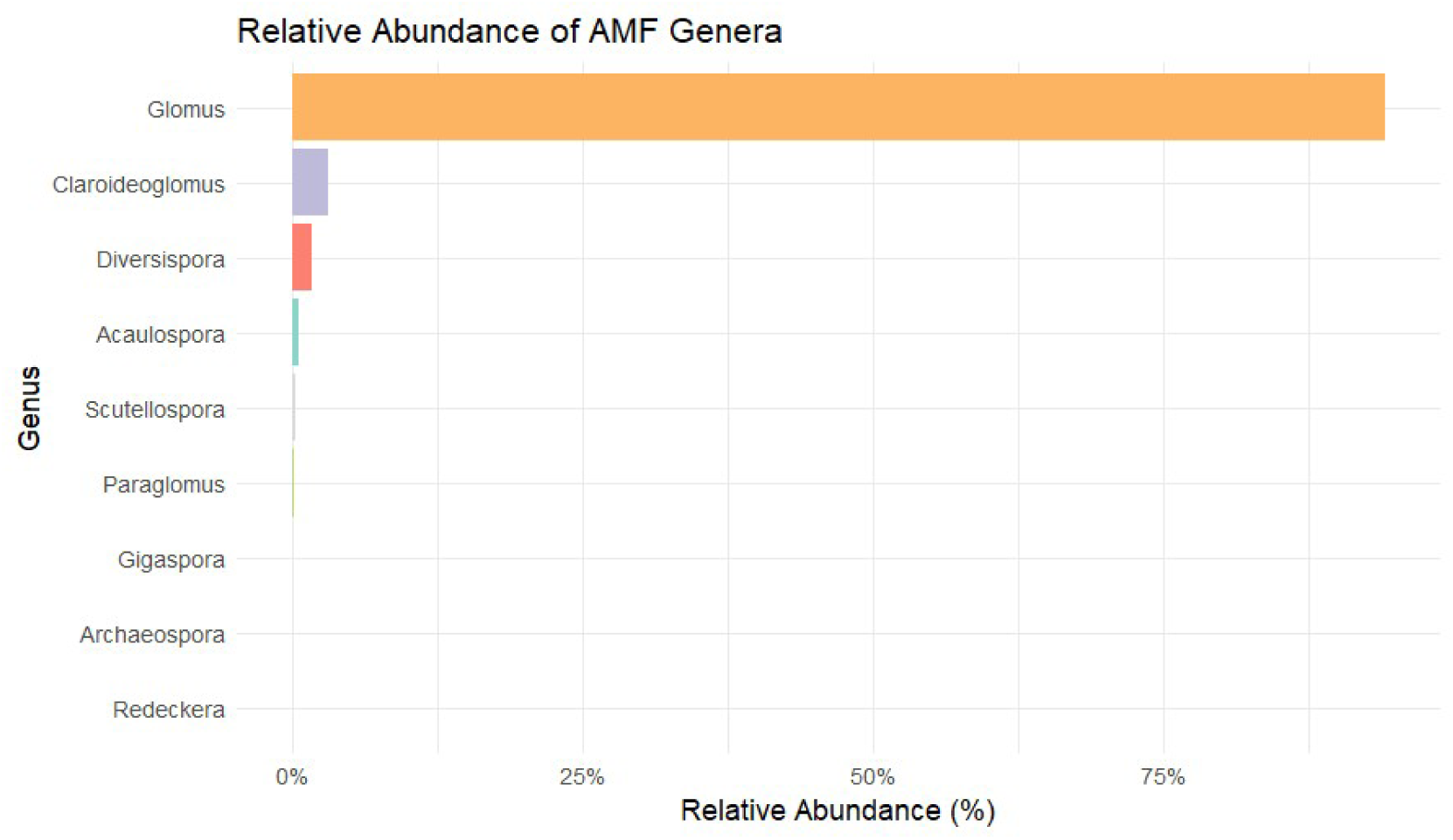
Relative abundance of arbuscular mycorrhizal fungi (AMF) genera across all root samples.

**Figure 8:**
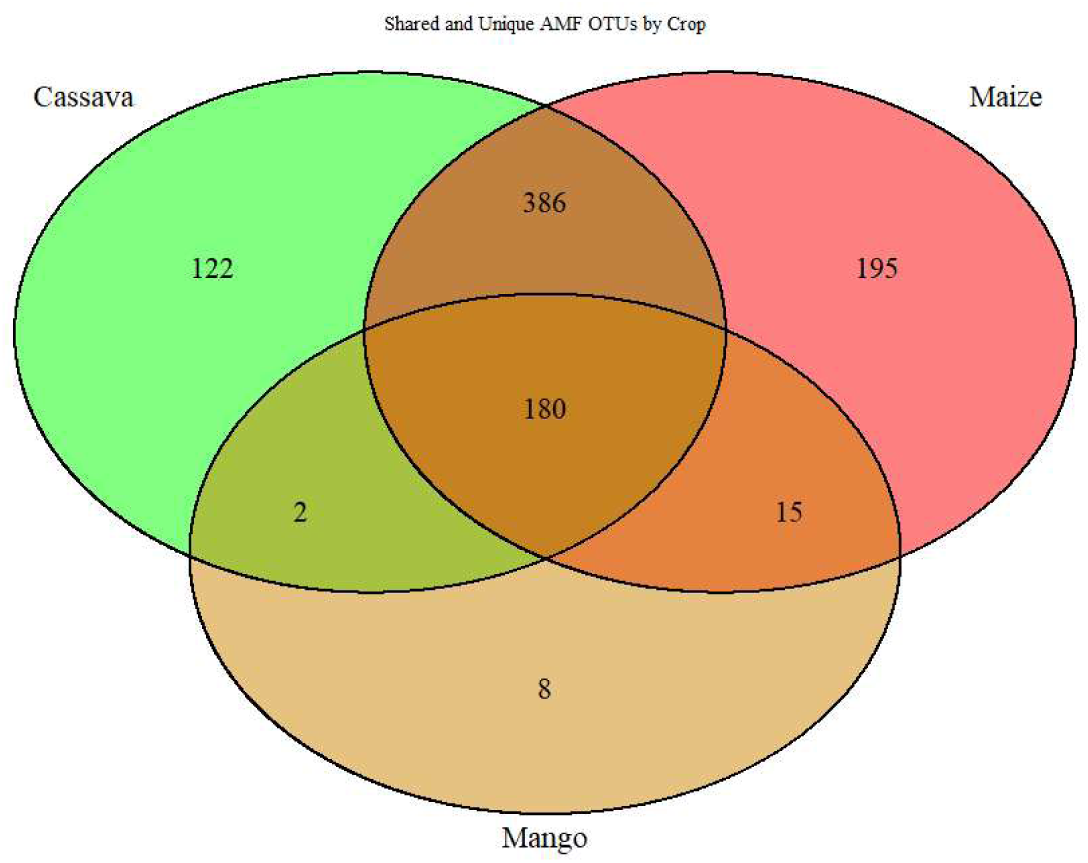
Shared and unique arbuscular mycorrhizal fungi (AMF) OTUs among cassava, maize, and mango roots.

**Figure 9:**
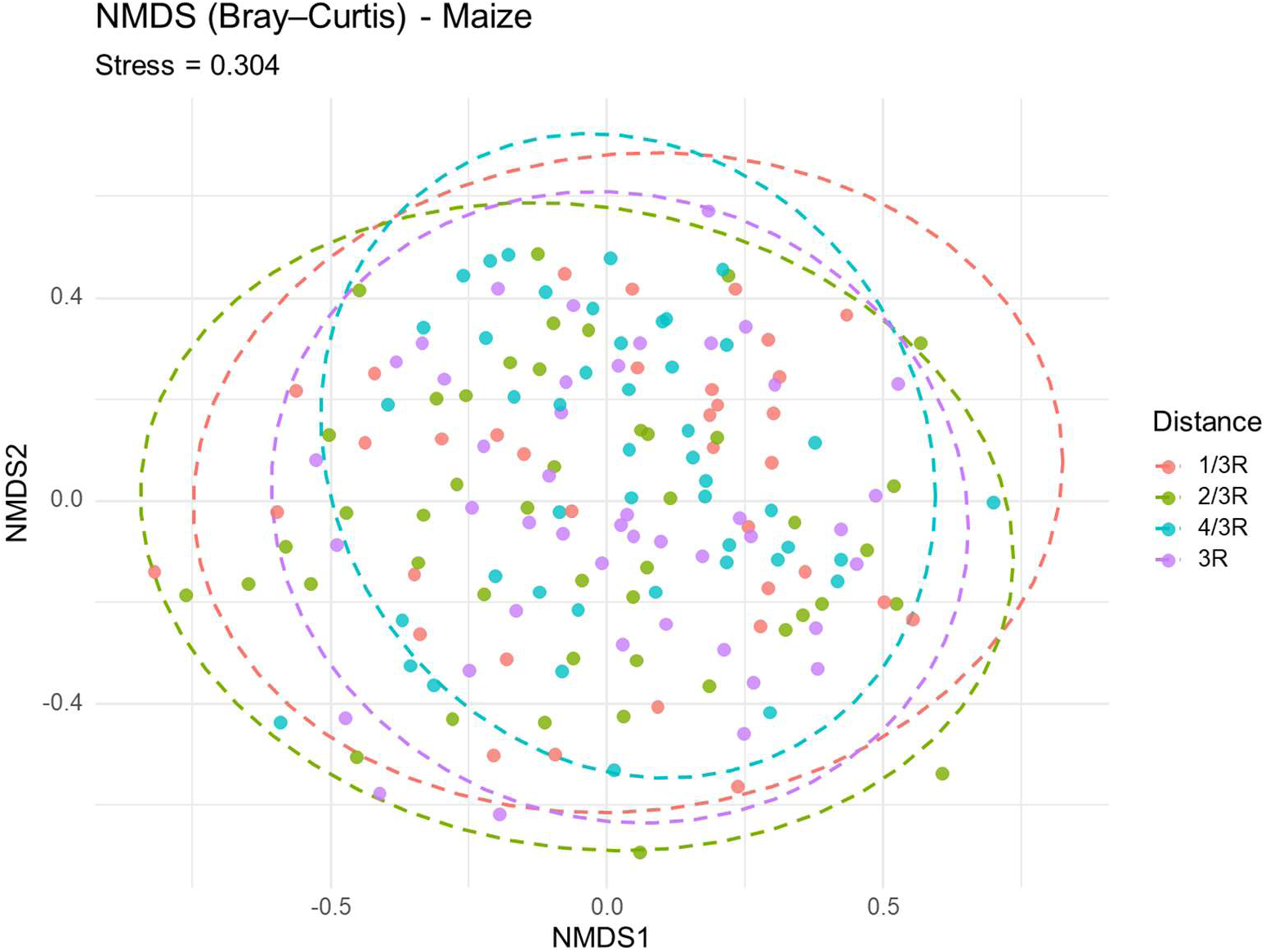
NMDS ordination based on Bray–Curtis dissimilarities showing AMF community composition in maize roots collected at four distances from Mangifera indica trees (1/3R, 2/3R, 4/3R, 3R). Overlapping ellipses reflect high community similarity across distances. Stress = 0.304.

**Figure 10:**
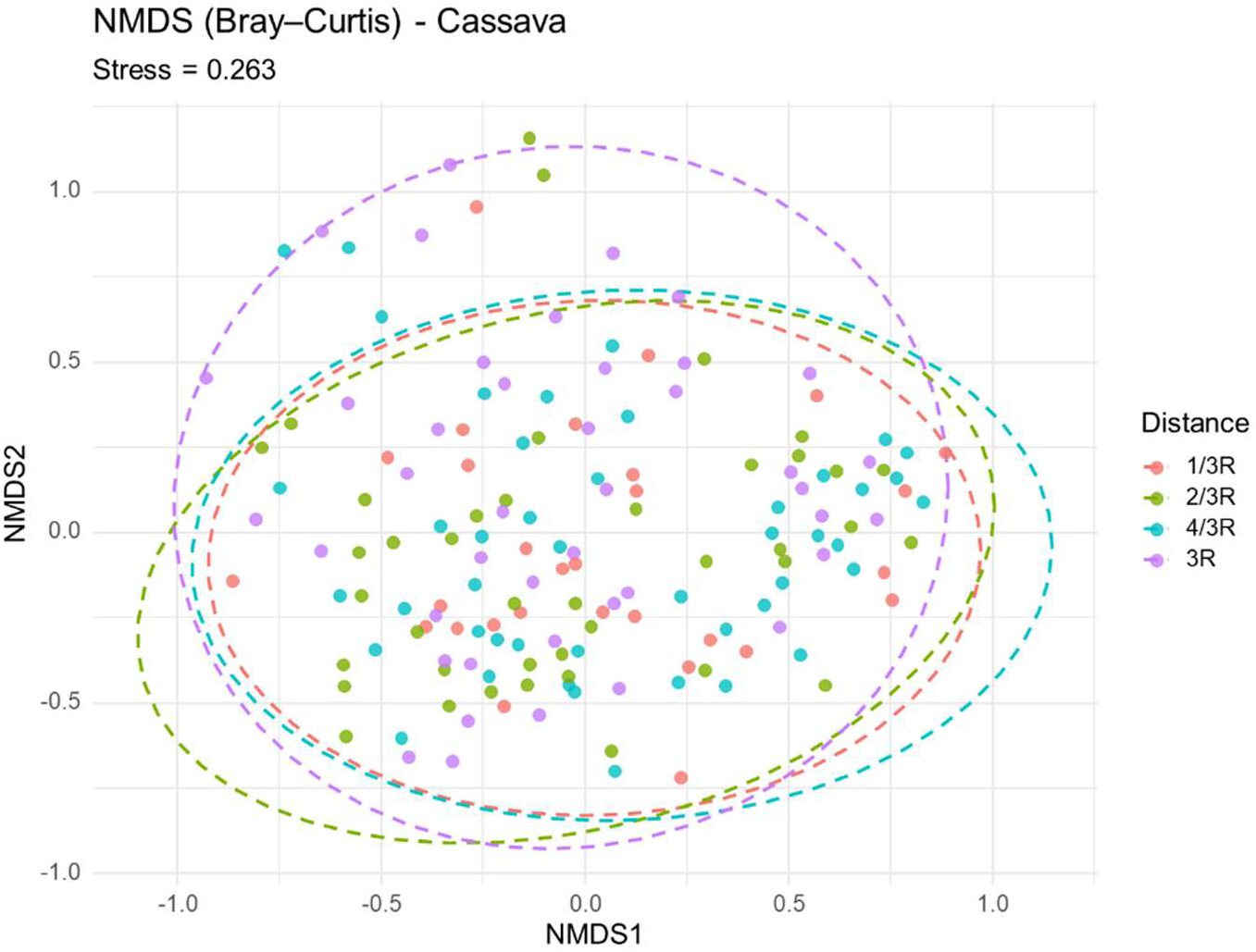
NMDS ordination based on Bray–Curtis dissimilarities showing AMF community composition in cassava roots sampled at four distances from Mangifera indica trees (1/3R, 2/3R, 4/3R, 3R). Overlapping ellipses indicate high similarity among distance classes. Stress = 0.263.

**Figure 6.**
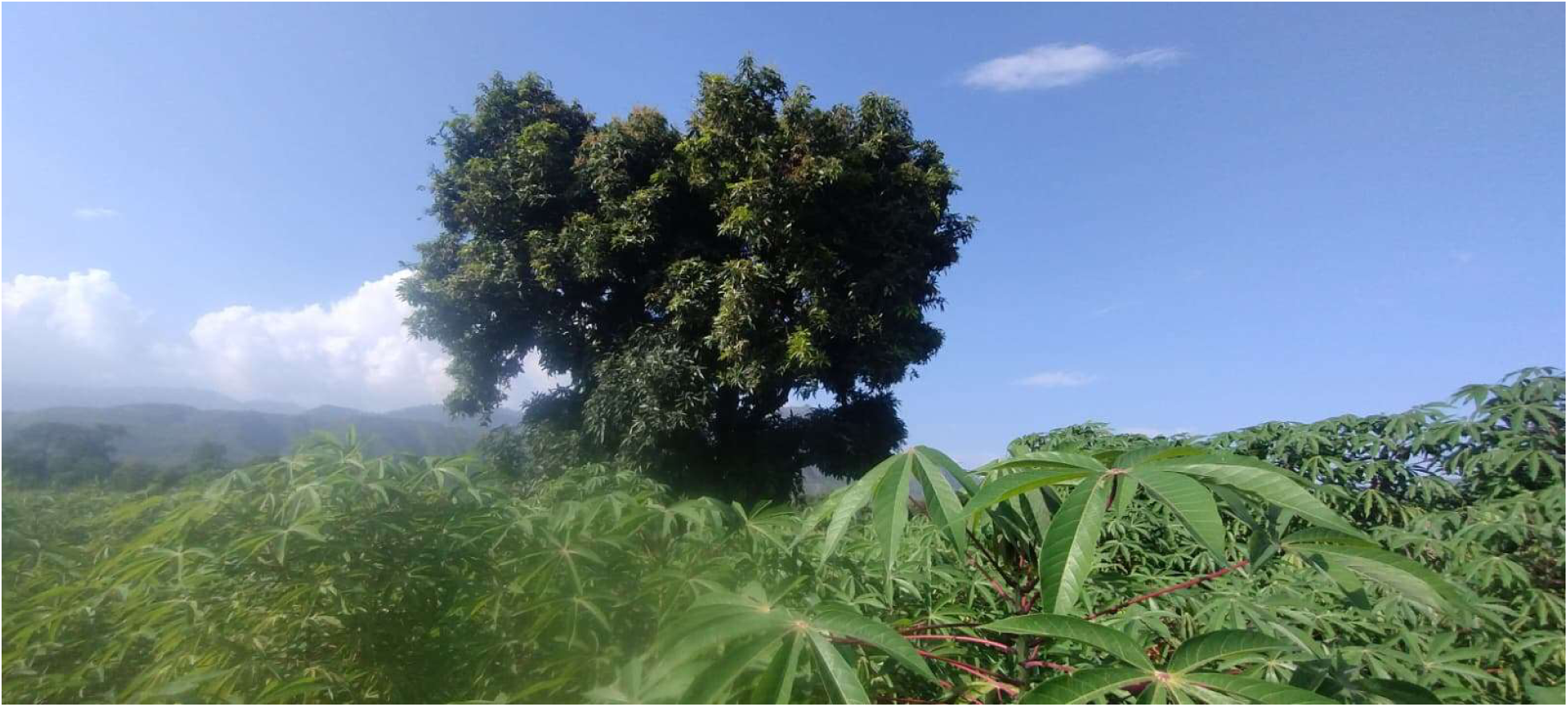
Field view of the cassava–mango system illustrating the sampling environment. Cassava plants are cultivated under mango canopy influence, where root and soil samples were collected along distance transects (1/3R, 2/3R, 4/3R, 3R) for AMF diversity assessment.

**Table 1:** Summary of statistical results testing the effect of distance from Mangifera indica trees on soil properties.

| Soil Properties | Effect | Df | F_value | p_value |
| --- | --- | --- | --- | --- |
| PH | Distance | 3, 330 | 3.74 | 0.01 |
| EC (dS/m) | Distance | 3, 330 | 32.7 | < 0.001 |
| Olsen. P(mg/kg) | Distance | 3, 330 | 1.37 | 0.25 |
| TN% | Distance | 3, 330 | 26.37 | < 0.001 |
| OC% | Distance | 3, 330 | 57.46 | < 0.001 |
| K<sub>av</sub>(mg/kg) | Distance | 3, 330 | 117.05 | < 0.001 |

**Table 2:**
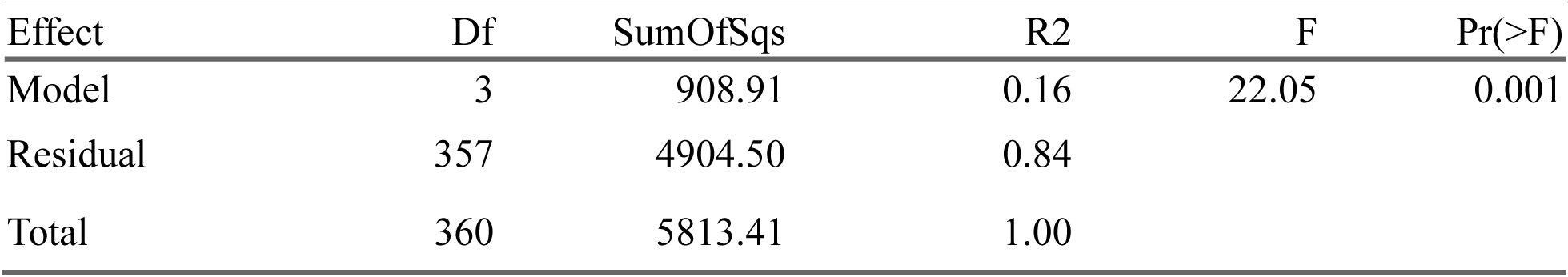
Results of permutational multivariate analysis of variance (PERMANOVA) testing the effect of distance from the tree base on the multivariate composition of soil properties in mango-based agroforestry systems.

| Effect | Df | SumOfSqs | R2 | F | Pr(>F) |
| --- | --- | --- | --- | --- | --- |
| Model | 3 | 908.91 | 0.16 | 22.05 | 0.001 |
| Residual | 357 | 4904.50 | 0.84 |  |  |
| Total | 360 | 5813.41 | 1.00 |  |  |

**Table 3:** Relative Abundance (%) of Indicator AMF Families among Hosts.

| *Family* | **Cassava** | **Maize** | **Mango** |
| --- | --- | --- | --- |
| *Acaulosporaceae* | 0.18 | 0.54 | 1.53 |
| *Claroideoglomeraceae* | 0.05 | 0.20 | 0.56 |
| *Diversisporaceae* | 2.55 | 1.80 | 0.58 |
| *Gigasporaceae* | 0.21 | 0.57 | 0.00 |
| *Glomeraceae* | 0.95 | 0.96 | 0.96 |
| *Paraglomeraceae* | 0.04 | 0.02 | 0.57 |

**Table 4:** Alpha-diversity of arbuscular mycorrhizal fungi (AMF) across Host types. Values are model estimates from Generalized Linear Mixed Models (GLMM), followed by post-hoc comparisons to assess differences among crops.

| Metric | Maize | Cassava | Mango | Test-statistic ($\chi^2$) | GLMM p-value | Post hoc p-value |
| --- | --- | --- | --- | --- | --- | --- |
| Sobs | 48.73 ± 1.84<sup>b</sup> | 54.12 ± 2.05<sup>c</sup> | 28.75 ± 2.00<sup>a</sup> | 72.5193 | < 0.001 | < 0.001 |
| Sexp | 128.93 ± 9.59<sup>b</sup> | 156.69 ± 11.66<sup>b</sup> | 69.64 ± 8.52<sup>a</sup> | 36.5344 | < 0.001 | < 0.001 |
| N1 | 9.08 ± 0.43<sup>b</sup> | 8.15 ± 0.44<sup>b</sup> | 4.95 ± 0.78<sup>a</sup> | 24.5173 | < 0.001 | < 0.001 |
| N2 | 5.56 ± 0.28<sup>b</sup> | 4.87 ± 0.28<sup>b</sup> | 3.14 ± 0.51<sup>a</sup> | 19.5219 | < 0.001 | < 0.001 |

**Table 5:** PERMANOVA results assessing distance-related variation in AMF community composition in the maize–mango system. Values represent F-statistics, R², and p-values based on 999 permutations.

| Effect | Df | SumOfSqs | R2 | F | Pr(>F) |
| --- | --- | --- | --- | --- | --- |
| Model | 3 | 0.848 | 0.016 | 0.893 | 0.662 |
| Residual | 166 | 52.568 | 0.984 |  |  |
| Total | 169 | 53.416 | 1.000 |  |  |

**Table 6:**
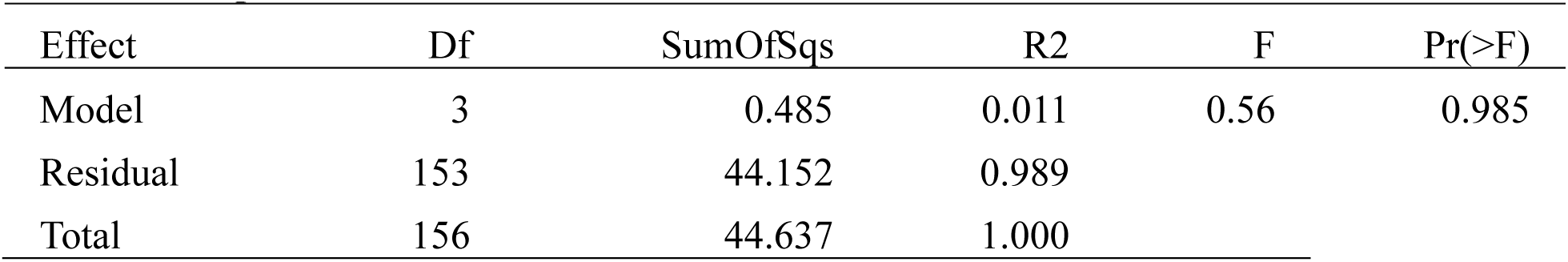
PERMANOVA results assessing distance-related variation in AMF community composition in the Cassava_mango systems. Values represent F-statistics, R², and p-values based on 999 permutations.

| Effect | Df | SumOfSqs | R2 | F | Pr(>F) |
| --- | --- | --- | --- | --- | --- |
| Model | 3 | 0.485 | 0.011 | 0.56 | 0.985 |
| Residual | 153 | 44.152 | 0.989 |  |  |
| Total | 156 | 44.637 | 1.000 |  |  |

**Table 7:** Results of permutational multivariate analysis of variance (PERMANOVA) testing the effect of Host root type on AMF composition in mango-based agroforestry systems.

| Effect | Df | SumOfSqs | R2 | F | Pr(>F) |
| --- | --- | --- | --- | --- | --- |
| Model | 2 | 6.76 | 0.06 | 11.91 | 0.0001 |
| Residual | 353 | 100.16 | 0.94 |  |  |
| Total | 355 | 106.92 | 1 |  |  |

**Table 8.** Pairwise PERMANOVA comparison of AMF community composition among Host types.

| Group1 | Group2 | R2 | p_value | p_adj |
| --- | --- | --- | --- | --- |
| Maize | Cassava | 0.0063 | 0.0253 | 0.0253 |
| Maize | Mango | 0.0869 | 0.0001 | 0.00015 |
| Cassava | Mango | 0.111 | 0.0001 | 0.00015 |

**Table 9:** AMF Diversity Response to Soil Properties in maize–Mango and Cassava_mango systems.

| Agroforestry Systems | Diversity Index | Soil Variable | Estimate | Std. Error | z-value | p-value | Pseudo R<sup>2</sup> |
| --- | --- | --- | --- | --- | --- | --- | --- |
| **Maize–Mango** | Sobs | K | -0.050 | 0.034 | -1.48 | 0.142 | 0.031 |
|  | Sobs | N | 0.083 | 0.034 | 2.46 | **0.015** | 0.031 |
|  | Sexp | O.C | 0.085 | 0.057 | 1.50 | 0.136 | 0.016 |
|  | N1 | N | 0.084 | 0.041 | 2.04 | **0.043** | 0.034 |
|  | N1 | P | -0.085 | 0.041 | -2.07 | **0.040** | 0.034 |
|  | N2 | N | 0.079 | 0.044 | 1.79 | **0.076** | 0.028 |
|  | N2 | P | -0.077 | 0.044 | -1.74 | **0.084** | 0.028 |
| **Cassava–Mango** | Sobs | N | 0.050 | 0.024 | 2.06 | **0.041** | 0.026 |
|  | Sexp | E.C | 0.079 | 0.060 | 1.31 | 0.194 | 0.079 |
|  | Sexp | P | 0.111 | 0.060 | 1.83 | **0.069** | 0.079 |

